# Topoisomerase IIα Prevents, But Does Not Resolve, Ultrafine Anaphase Bridges By Two Mechanisms

**DOI:** 10.1101/815001

**Authors:** Simon Gemble, Géraldine Buhagiar-Labarchède, Rosine Onclercq-Delic, Sarah Lambert, Mounira Amor-Guéret

## Abstract

Topoisomerase IIα (Topo IIα), a well-conserved double-stranded DNA (dsDNA)-specific decatenase, processes dsDNA catenanes resulting from DNA replication during mitosis. Topo IIα defects lead to an accumulation of ultrafine anaphase bridges (UFBs), a type of chromosome non-disjunction. Topo IIα has been reported to resolve DNA anaphase threads, possibly accounting for the increase in UFB frequency upon Topo IIα inhibition. We hypothesized that the excess UFBs might also result, at least in part, from an impairment of the prevention of UFB formation by Topo IIα. We found that Topo IIα inhibition promotes UFB formation without affecting UFB resolution during anaphase. Moreover, we showed that Topo IIα inhibition promotes the formation of two types of UFBs depending on cell-cycle phase. Topo IIα inhibition during S-phase compromises complete DNA replication, leading to the formation of UFB-containing unreplicated DNA, whereas Topo IIα inhibition during mitosis impedes DNA decatenation at metaphase-anaphase transition, leading to the formation of UFB-containing DNA catenanes. Thus, Topo IIα activity is essential to prevent UFB formation in a cell-cycle dependent manner, but dispensable for UFB resolution during anaphase.

## Introduction

Genome stability requires accurate DNA replication during S-phase and on correct chromosome segregation during mitosis. Errors impairing these two crucial steps are particularly prone to induce genetic instability [1, 2]. DNA replication leads to the formation of intertwines between two DNA strands, referred to as DNA catenanes, the resolution of which requires the introduction of transitory breaks. Topoisomerases play a key role in DNA catenane processing. Topoisomerase IIα (Topo IIα) is a well-conserved double-stranded DNA (dsDNA)-specific decatenase enzyme that processes dsDNA catenanes [2-5]. Topo IIα activity leads to double strand breakage followed by intra-molecular strand passage and DNA re-ligation [2]. The decatenating activity of Topo IIα plays a major role in several aspects of chromosome dynamics, including DNA replication and chromosome segregation [6].

Topoisomerase activity ahead of the replication fork cannot resolve all dsDNA catenanes. Moreover, convergence of two replisomes leads to the steric hindrance of topoisomerase activity [2, 7]. Consequently, some dsDNA catenanes are not resolved before the onset of mitosis. They form physical links between the sister chromatids and must therefore be processed by Topo IIα before chromosome segregation in anaphase [8]. Indeed, the disruption of Topo IIα activity leads to incomplete sister chromatid disjunction [9, 10]. Sister-chromatid anaphase bridges are of two types: chromatin anaphase bridges that can be stained with conventional dyes, such as DAPI, and ultrafine anaphase bridges (UFBs) that cannot be stained with conventional dyes or antibodies against histones. Both chromatin and ultrafine anaphase bridges result from a defect in sister chromatid segregation. During mitosis, PICH (Plk1-interacting checkpoint helicase), an SNF2-family DNA translocase involved in chromosome segregation [10-14], is recruited on both chromatin bridges and UFBs [10-14]. UFBs were discovered in 2007 [11, 13], and are present in all cell lines tested and are, thus, considered to be physiological structures [13, 15]. Most UFBs are of centromeric origin, but some originate from common fragile sites, telomeres or ribosomal DNA repeats [13, 14, 16, 17]. UFBs were reported to contain either unresolved DNA catenations, or replication intermediates [11-13, 15, 18]. In a previous study, we reported that the intracellular accumulation of dCTP, due to cytidine deaminase (CDA) deficiency, leads to an excess of UFB-containing unreplicated DNA, due to a decrease in the basal activity of poly(ADP-ribose) polymerase 1 (PARP-1), which promotes the premature entry of cells into mitosis, before completion of DNA replication has been completed [15, 18].

Topo IIα inhibition leads to a large increase in the frequency of centromeric UFBs [10, 11, 13, 19-21]. In this study, we investigated the molecular origin of the increase in UFB frequency following Topo IIα inhibition. We found that Topo IIα inhibition led to two types of UFBs, the type of UFB formed depending on the phase of the cell cycle. Topo IIα inhibition during S-phase impairs DNA replication, leading to the formation of UFB-containing unreplicated DNA during mitosis, whereas Topo IIα inhibition during mitosis prevents DNA decatenation, resulting in UFB-containing dsDNA catenanes. Thus, Topo IIα inhibition impairs both DNA replication during S-phase and DNA decatenation during mitosis, leading to the formation of two types of UFB with different molecular origins. More importantly, we also found that Topo IIα inhibition had no effect on the kinetics of UFB resolution during the progression of mitosis, excluding a role for Topo IIα activity in UFB resolution. Our results therefore demonstrate that Topo IIα activity is required to prevent the formation of UFBs through replication defects or a lack of resolution of DNA catenanes when cells enter mitosis, but that this enzyme is not involved in the resolution of pre-existing UFBs during anaphase.

## Results & Discussion

### Topoisomerase IIα activity is dispensable for UFB resolution during mitosis

Topo IIα inhibition leads to a large increase in UFB frequency, and thus it has been proposed that Topo IIα activity is required for UFB resolution, accounting for the increase in UFB frequency upon Topo IIα inhibition [10, 11, 13, 19-21]. However, the increase in UFB frequency upon Topo IIα inhibition could also reflect, at least in part, an accumulation of newly formed UFB. We therefore first investigated whether Topo IIα inhibition compromised the resolution or the formation of UFBs.

Topo IIα inhibition in HeLa cells with ICRF-159 (1 or 10 μM, 8 h), a catalytic Topo IIα specific inhibitor [23], led to a dose-dependent increase in UFB frequency in anaphase cells in a dose-dependent manner, as expected (Figure 1A-C). We investigated whether Topo IIα inhibition affected UFB formation or resolution, by treating cells with ICRF-159 from S-phase until the end of mitosis and quantifying UFBs during mitosis: from metaphase (the first step in mitosis, during which the distance between sister chromatids is sufficiently large for the visualization of UFBs) to telophase. Using this approach, we were able to assess UFB formation (by determining the increase in UFB frequency over time) and the kinetics of UFB resolution (visualized as a decrease in UFB frequency during mitosis) (Figure 1D).

**Figure 1:**
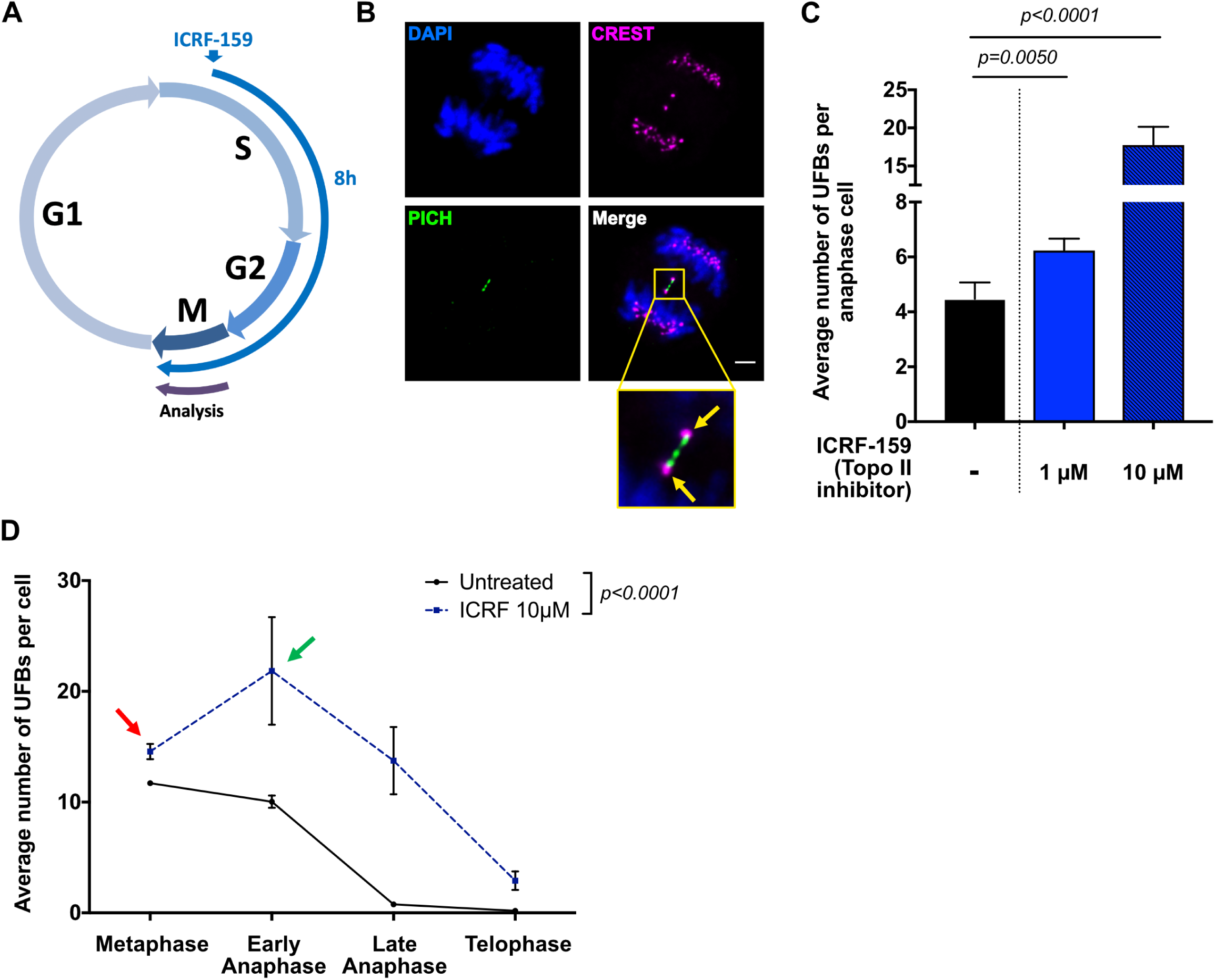
Topoisomerase IIα is not involved in UFB resolution. (A) Schematic representation of 8 hours of Topo IIα inhibition during the cell cycle; only cells treated during S-phase to mitosis were analyzed in anaphase. (B) Representative immunofluorescence deconvoluted *z*-projection images of PICH-positive UFBs in HeLa anaphase cells. DNA was visualized by DAPI staining (blue). Centromeres were stained with CREST serum (in magenta) and UFBs were stained with PICH antibody (in green). In the enlarged image, CREST foci at the extremities of the UFB are indicated by yellow arrows. Scale bar: 5 μm. (C) Bar graph presenting the mean number of UFBs per anaphase cell in HeLa cells, for cells left untreated (black bar) or treated with 1 or 10 µM ICRF-159 for 8 hours (blue bars); Errors bars represent means ± SD from three independent experiments (50-100 anaphase cells analysed per condition). (D) Mean number of UFBs per mitotic cells, from metaphase to telophase, for cells left untreated (continuous line) or treated with 10 µM ICRF-159 for 8 hours (discontinuous line); n=3, > 150 mitotic cells analyzed per condition. Statistical significance was assessed in *t*-tests (C) or by two-way ANOVA (D).

ICRF-159 treatment led to an increase in UFB frequency at metaphase (red arrow, Figure 1D). Thus, cells entered mitosis with with a higher frequency of UFBs when Topo IIα was inhibited. Interestingly, the frequency of UFBs was also much higher at the metaphase-anaphase transition (green arrow, Figure 1D), reflecting the formation of new UFBs early in mitosis. UFB frequency then decreased over time until the end of mitosis, in a similar manner in both untreated and ICRF-159-treated cells. UFBs are therefore resolved even if Topo IIα is inhibited.

Our data indicate that Topo IIα activity is dispensable for the resolution of pre-existing UFBs during mitosis, but is strictly necessary to prevent the formation of new UFBs. Our observations also indicate that Topo IIα inhibition promotes UFB formation in two different ways: before the onset of mitosis, as revealed by the increase in UFB frequency at metaphase, and during mitosis, leading to an increase in UFB frequency at the metaphase-anaphase transition.

### Topoisomerase IIα inhibition impairs complete DNA replication

We previously reported that delaying entry into mitosis allows the completion of DNA replication and prevents the formation of UFBs, strongly suggesting that, in unchallenging condition, these structures result from the accumulation of unreplicated DNA during mitosis [15, 18]. Topo IIα inhibition leads to an increase in UFB frequency at metaphase (Figure 1D). We therefore first investigated whether Topo IIα inhibition prevented the completion of DNA replication, leading to the formation of new UFB-containing unreplicated DNA on entry into mitosis.

We therefore determined whether centromere replication was impaired upon Topo IIα inhibition. Cells were left untreated or were treated with ICRF-159 for 8 h. We used CREST staining to quantify double-dotted (yellow arrows, Figure 2A) and single-dotted (white arrow, Figure 2A) foci in prometaphase corresponding to fully replicated and unreplicated centromeres, respectively (Figure 2A), as previously described (Gemble et al., 2015). The frequency of unreplicated centromeres was significantly higher in cells treated with 1 or 10 µM ICRF-159 for 8 hours than in control cells (Figure 2A and B), demonstrating that Topo IIα inhibition impaired the replication of centromeric DNA. Interestingly, the frequency of unreplicated centromeres did not differ between cells treated with 1 and 10 µM ICRF-159, contrasting with the dose-dependent effect of ICRF-159 on UFB formation (Figure 1C).

**Figure 2:**
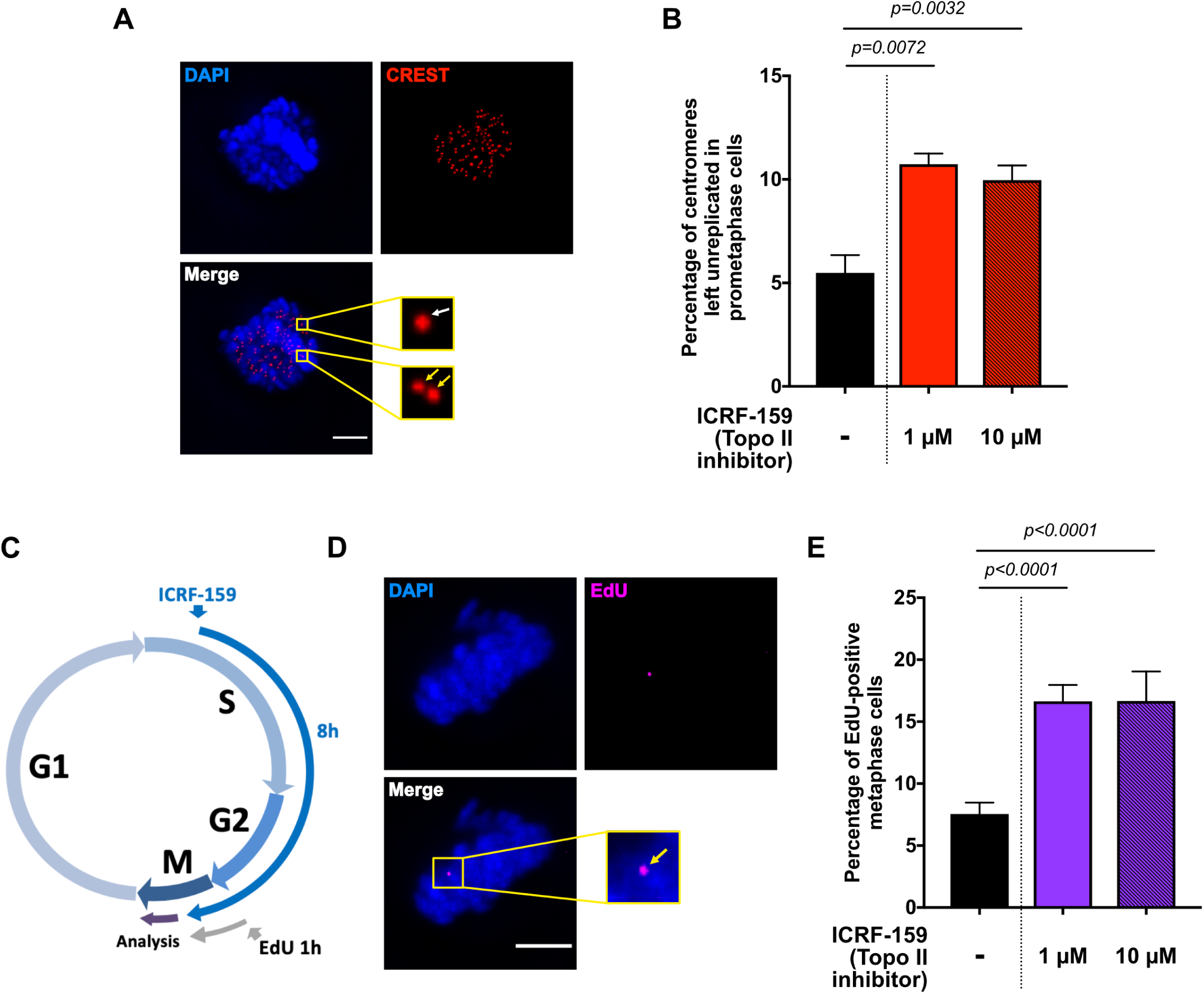
Topoisomerase IIα inhibition impairs complete DNA replication. (A) Representative immunofluorescence deconvoluted *z*-projection images of a prometaphase HeLa cell. DNA was visualized by DAPI staining (blue). Centromeres were stained with CREST serum (in red). Boxed images are enlarged; single-dotted CREST foci are indicated by white arrows and double-dotted CREST foci are indicated by yellow arrows. Scale bar: 5 µm. (B) Bar graph showing the percentage of centromeres left unreplicated in HeLa prometaphase cells left untreated (black bar) or treated with 1 or 10 µM ICRF159 for 8 hours (red bars). Error bars represent means ± SD from three independent experiments (> 90 prometaphase cells per condition). (C) Schematic representation of 8 hours of Topo IIα inhibition during the cell cycle. Only cells treated during S-phase to mitosis were analyzed in anaphase. EdU was added one hour before analysis. (D) Representative immunofluorescence deconvoluted *z*-projection images of a metaphase HeLa cell with EdU incorporation. DNA was visualized by DAPI staining (blue). EdU was stained with Alexa Fluor 555 (in magenta). Enlarged image shows one EdU focus on mitotic chromosomes (yellow arrow). Scale bar: 5 µm. (E) Bar graph presenting the percentage of HeLa metaphase cells presenting EdU foci after being left untreated (black bar) or after treatment with 1 or 10 µM ICRF159 for 8 hours (purple bars) and presenting EdU foci. Error bars represent means ± SD for three independent experiments (100-200 metaphase cells per condition were analyzed). Statistical significance was assessed in *t*-test.

For confirmation of the effect of Topo IIα inhibition on DNA replication, we then evaluated the levels of mitotic DNA synthesis (MiDAS). MiDAS contributes to the processing of unreplicated DNA sequences during mitosis and can therefore be used to detect problems leading to incomplete DNA replication during the previous S-phase [15, 18, 24-26]. MiDAS can be visualized by 5-ethynyl-2’-deoxyuridine (EdU) incorporation, leading to the formation of foci on condensed chromosomes (yellow arrow, Figure 2D). We found that treatment with 1 or 10 µM ICRF-159 treatment for 8 hours led to a significant increase in the percentage of prometaphase cells presenting MiDAS, with no dose dependence (Figure 2C-E). These data confirm that Topo IIα inhibition results in an accumulation of unreplicated DNA during mitosis, reflecting incomplete DNA replication in the previous S-phase. These observations are consistent with several studies in yeast or *in vitro*, showing that Topo IIα facilitates DNA replication [2, 27-30]. They also suggest that Topo IIα activity is essential to promote complete DNA replication in mammalian cells.

Our data demonstrate that Topo IIα inhibition impairs the completion of DNA replication, probably leading to the formation of UFB-containing unreplicated DNA on entry into mitosis.

### Topoisomerase IIα inhibition promotes the formation of two different types of UFBs depending on the phase of the cell cycle

Our findings that Topo IIα inhibition increases total UFB frequency in a dose-dependent manner, but impairs DNA replication independently of ICRF-159 concentration, suggest that Topo IIα inhibition affects another process of UFB formation, in addition to DNA replication. Topo IIα activity is required for both the completion of DNA replication during S-phase (Figure 2 and [2, 27-30]) and DNA decatenation during mitosis [8]. We therefore investigated the respective contributions of these processes to the increase in UFB formation in response to Topo IIα inhibition. We analyzed UFB frequency in HeLa anaphase cells after treatment either during S-phase (addition of ICRF-159 for 6 hours followed by a release period of 4 hours), or during mitosis (addition of ICRF-159 1 hour before UFB analysis) in anaphase cells (Figure 3A). We confirmed the cell cycle phase-specificity of our treatment by simultaneously treating cells only during S-phase or only during mitosis (Figure 3A), with both ICRF-159 and EdU. As expected, all cells treated with ICRF-159 during S-phase and then analyzed during mitosis were positive for EdU (Figure 3B), whereas cells treated only during mitosis were EdU-negative (Figure 3B). Consistent with these results, we found that Topo IIα inhibition during S-phase led to an increase in the percentage of unreplicated centromeres during mitosis and to an increase in the level of MiDAS (Figure 3C and D), whereas inhibition during mitosis did not, confirming our previous findings (Figure 2) and demonstrating the cell cycle specificity of the treatment. These results indicate that Topo IIα activity is required during S-phase, to promote complete DNA replication.

**Figure 3:**
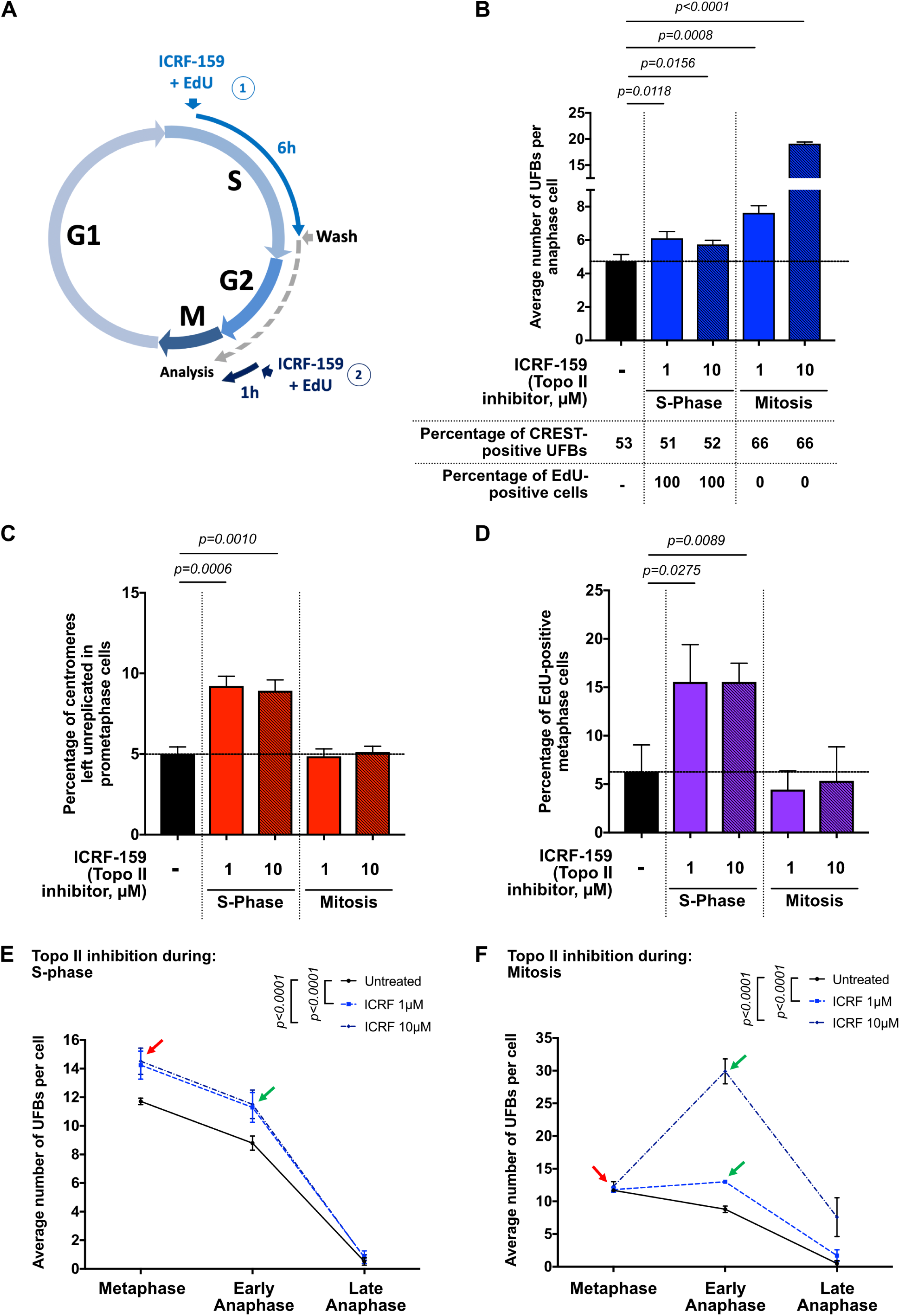
Topoisomerase IIα inhibition promotes two different types of UFB, depending of the phase of the cell cycle. (A) Schematic representation of Topo IIα inhibition during the cell cycle; only cells treated during S-phase and then released (1) or treated during mitosis (2) were analyzed in anaphase. Percentages of CREST-positive UFBs and EdU-positive cells for each condition are indicated below the graph. (B) Bar graph showing the mean number of UFBs per anaphase cell in HeLa cells left untreated (black bar) or treated with 1 or 10 µM ICRF-159 during S-phase or during Mitosis (blue bars); Errors bars represent means ± SD from three independent experiments (> 85 anaphase cells analysed per condition). (C) Percentage of centromeres left unreplicated in HeLa prometaphase cells left untreated (black bar) or treated with 1 or 10 µM ICRF159 during S-phase or mitosis (red bars). Error bars represent means ± SD from three independent experiments (> 75 prometaphase cells per condition). (D) Percentage of HeLa metaphase cells presenting EdU foci after being left untreated (black bar) or after treatment with 1 or 10 µM ICRF159 during S-phase or mitosis (purple bars). Error bars represent means ± SD for three independent experiments (> 90 metaphase cells per condition were analysed). (E) Bar graph showing the mean number of UFBs per mitotic cells, from metaphase to anaphase, for cells left untreated (continuous line) or treated with 1 or 10 µM ICRF-159 during S-phase (discontinuous lines; n=5, 90-165 mitotic cells analysed per condition). (F) Mean number of UFBs per mitotic cells, from metaphase to anaphase, for cells left untreated (continuous line) or treated with 1 or 10 µM ICRF-159 during mitosis (discontinuous lines; n=5, 90-165 mitotic cells analyzed per condition). Statistical significance was assessed with *t*-test (B; C and D) or by two-way ANOVA test (E and F).

In the same experimental conditions, Topo IIα inhibition during S-phase led to a slight, but significant, dose-independent increase in the mean number of UFBs per cell (Figure 3B), probably due to the effect of Topo IIα inhibition on the completion of DNA replication (Figure 2 and Figure 3C-D). However, in cells treated with Topo IIα inhibitor only during mitosis, we observed a much higher dose-dependent increase in UFB frequency (Figure 3B). These data suggest that most of the UFBs observed upon Topo IIα inhibition result from the loss of Topo IIα activity during mitosis. Moreover, our observations suggest that most of the UFBs-containing unreplicated DNA (resulting from Topo IIα inhibition during S-phase) are probably resolved before entry in mitosis.

We then investigated the effect of Topo IIα inhibition on the formation of different types of UFB as a function of the phase of the cell cycle. We treated cells with ICRF-159 (1 and 10 μM) during S-phase or during mitosis, and we analyzed UFB frequency in mitotic cells, from metaphase to anaphase (Figure 3E and F). Topo IIα inhibition during S-phase led to an increase in the mean number of UFBs per cell at metaphase (red arrow, Figure 3E), indicating that the cells entered mitosis with more UFBs. However, Topo IIα inhibition during S-phase was not associated with the formation of new UFBs at metaphase-anaphase transition (green arrow, Figure 3E). UFB frequency decreased during the course of mitosis, with kinetics similar to those observed in untreated cells, consistent with the resolution of UFBs independently of Topo IIα activity. We therefore hypothesized that Topo IIα inhibition in S-phase would impair complete DNA replication, leading to the formation of UFB-containing unreplicated DNA on entry into mitosis. By contrast, the restriction of Topo IIα inhibition to mitosis had no effect on UFB frequency at metaphase (red arrow, Figure 3F). However, UFB frequency was much higher at the metaphase-anaphase transition, particularly in response to 10 μM ICRF-159, (green arrow, Figure 3F), reflecting the formation of new UFBs during mitosis. UFB frequency subsequently decreased during anaphase, confirming that Topo IIα activity was dispensable for UFB resolution (Figure 3F). Interestingly, the increase in UFB frequency at metaphase-anaphase transition was not observed in cells treated only during S-phase (green arrow, Figure 3E). Topo IIα activity is required to resolve centromeric DNA catenations at the metaphase-anaphase transition [6]. We therefore suggest that DNA decatenation is compromised when Topo IIα is inhibited during mitosis, promoting the formation of UFB-containing DNA catenanes in anaphase. Consistent with this hypothesis, we observed that most of the UFBs generated by Topo IIα inhibition during mitosis were of centromeric origin (Figure 3B). This region of the chromosome has already been shown to be associated with UFB-containing DNA catenanes [10, 13, 20, 21]. More importantly, UFB frequency decreased from early anaphase to late anaphase in cells treated with ICRF-159 during S-phase, but also in those treated during mitosis, indicating that UFBs were resolved in both cases, despite the maintenance of Topo IIα inhibition during mitosis. It has been previously shown that Topo IIα inhibition in chemically-synchronized mitotic cells results in the accumulation of PICH-positive threads, suggesting that Topo IIα activity is necessary to resolve these structures [19]. However, the specific fate of UFBs was not addressed. Here, by maintaining Topo IIα inhibition during mitosis and by specifically analyzing UFB frequency from metaphase to late anaphase, we clearly demonstrated that Topo IIα activity is dispensable for UFB resolution during mitosis.

### Topoisomerase IIα activity is dispensable for UFB resolution

It has been reported that aberrant UFB resolution at the end of mitosis causes DNA damage leading to the formation of 53BP1 bodies in the next G1 phase, to protect broken DNA ends until repair [31]. We investigated whether UFB resolution in cells treated with Topo IIα inhibitors was aberrant, by analyzing the number of 53BP1 bodies in cells left untreated or treated with ICRF-159 during S-phase or during mitosis (Figure 4A-C). We ensured that only cells entering G1 after Topo IIα inhibition were analyzed by treating cells with EdU before adding ICRF-159 during S-phase or mitosis (Figure 4A), and analyzing only the EdU-positive G1 cells. In both cases (inhibition during S phase and inhibition during mitosis), we observed a slight increase in the number of 53BP1 bodies in G1 cells (Figure 4B-C) potentially reflecting double-strand breaks generated during mitosis or under-replicated genomic loci converted to DNA or chromatin lesions and transmitted to daughter cells to be repaired, as previously reported [31]. However, despite the observed dose-dependent increase in UFB frequency in cells treated during mitosis with ICRF-159 during mitosis (Figure 3B), the number of 53BP1 foci in the following interphase was similar in cells treated with Topo IIα inhibitor at concentrations of 1 or 10 µM (Figure 4C). Moreover, Topo IIα inhibition led to a larger increase in UFB frequency when ICRF-159 was added during mitosis that when it was added during S-phase (Figure 3B), but we observed no significant difference in the number of 53BP1 foci in G1 between Topo IIα inhibition during S-phase and Topo IIα inhibition during mitosis. Together, these observations show that the occurrence of DNA damage in G1 was not coupled to UFB frequency in the previous mitosis, indicating that most of UFBs are resolved during anaphase when Topo IIα is inhibited. These results confirm that Topo IIα activity is dispensable for UFB resolution during mitosis.

**Figure 4:**
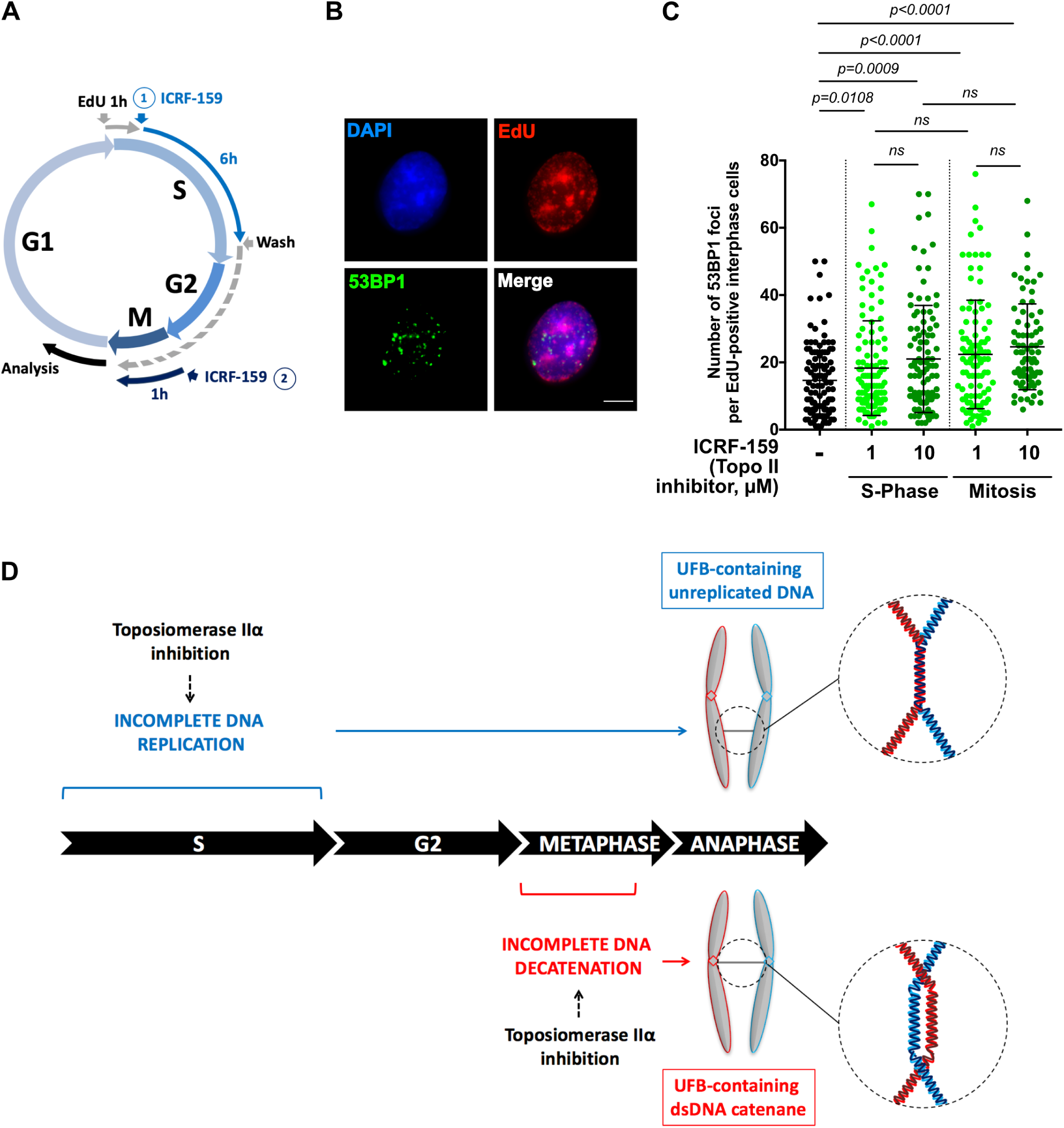
Topoisomerase IIα activity is dispensable for UFB resolution. (A) Schematic representation of Topo IIα inhibition during the cell cycle. Only cells treated during S-phase and then released (1) or treated during mitosis (2) were analyzed in G1. EdU was added for 1 hour during S-phase. Only EdU-positive cells were analyzed. (B) Representative immunofluorescence deconvoluted z-projection images of an interphase HeLa cell with EdU incorporation. DNA was visualized by DAPI staining (blue). EdU was stained with Alexa Fluor 555 (in red). DNA damage was detected by staining with the 53BP1 antibody (in green). Scale bar: 5 µm. (C) Dot blot presenting the number of 53BP1 foci per EdU-positive interphase cells in HeLa cells, for cells left untreated (black bar) or treated with 1 or 10 µM ICRF-159 during S-phase or during mitosis (green bars); Errors bars represent means ± SD from two independent experiments (> 100 interphase cells analyzed per condition). (D) Topo IIα inhibition leads to two types of UFBs, depending on the phase of the cell cycle. Topo IIα inhibition during S-phase compromises complete DNA replication, leading to the accumulation of unreplicated DNA in mitosis, resulting in an increase in the formation of UFB-containing unreplicated DNA. By contrast, Topo IIα inhibition during mitosis jeopardizes complete DNA decatenation process at the metaphase-anaphase transition, leading to the formation of UFB-containing DNA catenanes. Statistical significance was assessed in *t*-test.

Overall, our data shed light on the molecular origin of the supernumerary UFBs observed following Topo IIα inhibition, showing that they correspond to newly formed UFBs and not to unresolved pre-existing UFBs. We found that Topo IIα inhibition during S-phase compromised the completion of DNA replication, leading to the accumulation of unreplicated DNA during mitosis, probably leading to the formation of UFB-containing unreplicated DNA. We also found that the restriction of Topo IIα inhibition to mitosis resulted in a much higher frequency of UFBs at the metaphase-anaphase transition, particularly in the presence of 10 μM ICRF-159, reflecting the formation of new UFBs during mitosis. These UFBs probably result from impaired DNA decatenation at the metaphase-anaphase transition and correspond to newly formed UFB-containing DNA catenanes (Figure 4D). Our data demonstrate that maintaining Topo IIα inhibition during mitosis affects neither the kinetics of UFB resolution after metaphase-anaphase transition nor the accumulation of DNA damage in the next G1. We conclude that Topo IIα is largely dispensable for the resolution of UFBs.

In conclusion, our findings further extend the role of Topo IIα activity during the cell cycle, by showing that Topo IIα is required for complete DNA replication but dispensable for UFB resolution during mitosis.

## Materials & methods

### Cell culture and treatments

HeLa cells were cultured in DMEM supplemented with 10% FCS as previously described [15].

ICRF-159 (Razoxane) was provided by Sigma Aldrich (R8657) and was added to the cell culture medium at a final concentration of 1 or 10 µM.

All cells were routinely checked for mycoplasma infection.

### Immunofluorescence microscopy

Immunofluorescence staining and analysis were performed as previously described [15]. Primary and secondary antibodies were used at the following dilutions: rabbit anti-PICH antibody (1:150; H00054821-D01P from Abnova); human CREST antibody (1:100; 15-234-0001 from Antibodies Inc); mouse anti-53BP1 antibody (1/500; MAB3802 from Millipore); Alexa Fluor 633-conjugated goat anti-human antibody (1:500; A21091 from Life Technologies); Alexa Fluor 555-conjugated goat anti-rabbit (1:500; A21429 from Life Technologies). Cell images were acquired with a 3-D deconvolution imaging system consisting of a Leica DM RXA microscope equipped with a piezoelectric translator (PIFOC; PI) placed at the base of a 63x PlanApo N.A. 1.4 objective, and a CoolSNAP HQ interline CCD camera (Photometrics). Stacks of conventional fluorescence images were collected automatically at a Z-distance of 0.2 mm (Metamorph software; Molecular Devices). Images are presented as maximum intensity projections, generated with ImageJ software, from stacks deconvolved with an extension of Metamorph software [22].

### EdU staining

EdU incorporation into DNA was visualized with the Click-it EdU imaging kit (C10338 from Life Technologies), according to the manufacturer’s instructions. EdU was used at a concentration of 2 µM for the indicated time. Cells were incubated with the Click-it reaction cocktail for 15 minutes. Cell images were acquired with a 3-D deconvolution imaging system consisting of a Leica DM RXA microscope equipped with a piezoelectric translator (PIFOC; PI) placed at the base of a 63x PlanApo N.A. 1.4 objective, and a CoolSNAP HQ interline CCD camera (Photometrics). Stacks of conventional fluorescence images were collected automatically at a Z-distance of 0.2 mm (Metamorph software; Molecular Devices). Images are presented as maximum intensity projections generated with ImageJ software, from stacks deconvolved with an extension of Metamorph software.

### Statistical analysis

At least three independent experiments were carried out to generate each dataset and the statistical significance of differences was calculated with Student’s *t-*test or two-way ANOVA, as indicated in the figure legends.

## Acknowledgments

This work was supported by grants from the Institut Curie (PICSysBio), the *Centre National de la recherche Scientifique* (CNRS), the *Ligue contre le Cancer* (*Comité de l’Essonne*), the *Association pour la Recherche sur le Cancer* (ARC, SFI20121205645), the *Agence Nationale de la Recherche* (ANR-14-CE14-0004-01) and by a fellowship awarded to S.G by the *Ministère de l’Education, de l’Enseignement Supérieur et de la Recherche* and the ARC (DOC20140601310), and Institut Curie (PIC SysBio).

## Author contributions

SG performed the experiments, participated in the design of the experiments and data analysis, generated the figures and cowrote the manuscript. GBL and ROD performed experiments. SL contributed to data analysis and preparation of the manuscript. MA-G supervised the study, analyzed the data and cowrote the manuscript.

